# The AlphaFold Database Ages

**DOI:** 10.1101/2025.06.22.660930

**Authors:** Ifigenia Tsitsa, Anja Conev, Alessia David, Suhail A Islam, Michael J E Sternberg

## Abstract

The AlphaFold database provides 200M protein structures predicted by AlphaFold2 and released in 2022 from sequences in UniProt in April 2021. However, of the 20,504 full-length human structures in the AlphaFold database, 631 entries conflict with UniProt (version 2025_03 i.e. release 3 of 2025 published Jun 18, 2025); and there is a similar discrepancy for other species. This highlights how bioinformatics resources, as exemplified by the AlphaFold database can rapidly age. Time flies in bioinformatics.

## Text

The AlphaFold database (AlphaFoldDB) ^1^ is a major bioinformatics resource that provides predicted three-dimensional models for over 200 million protein sequences obtained from UniProt ^2^. These models were generated using AlphaFold2 ^3^, the deep learning algorithm developed by DeepMind. The importance of this database is underscored by its subsequent incorporation into UniProt ^2^, one of the most widely used protein databases. These models are extensively used by the community for tasks including molecular replacement (e.g. Phaser ^4^ and MolRep ^5^), missense variant prediction (e.g. FoldX ^6^ and Missense3D ^7^) and as templates in structure prediction (e.g. SwissModel ^8^ and Phyre2.2 ^9^). AlphaFoldDB was based on the protein sequences from April 2021 (UniProtKB version 2021_02, April 2021) which are outdated as UniProt sequences are regularly updated, a discrepancy that researchers should be aware of when studying specific proteins. Moreover, this is a major challenge for developers of general bioinformatics resources that require the maintenance of integrative and robust data management. Here, we have compared AlphaFold models with their respective UniProt sequences across six species and identified these discrepancies. It is vital for the community to be aware of these challenges to preserve the value of such resources.

In August 2025, we performed a comparison of AlphaFoldDB models with latest UniProt (version 2025_03 i.e. release 3 of 2025 published Jun 18, 2025)) for the human proteome revealed that, among 20,504 unique UniProt accessions with AlphaFold2-predicted structures, 631 sequences differed, corresponding to a 3.08% discrepancy between AlphaFold and UniProt sequences (Figure 1). Out of the conflicting 631 models, 116 correspond to the UniProt sequence entries which were made redundant since the AlphaFoldDB construction based on UniProtKB release 2021_02 ^10^. The redundant UniProt sequences are generally the result of obsolete or reassigned genes, updated taxonomic data, changes in reference databases, annotation errors, or periodic data consolidation to improve consistency and accuracy (*example: Q5TG92*). Sequence length mismatches were observed for 295 models, with UniProt sequences being longer than AlphaFold2 for 155 models and shorter for 140 models. For example, ZNT2 (Q9BRI3), a zinc transporter protein implicated in the transient neonatal zinc deficiency, has a region of approximately 50 amino acids inserted in the new UniProt sequence around position 100 of the sequence of the outdated AlphaFold. To assess the impact of this insertion on the ZNT2 structure, we compare model structures (technical details in *Supplementary Data*). In particular, we compared the AlphaFoldDB model (Figure 1B, blue) with an AlphaFold2 model generated from the updated UniProt sequence (Figure 1B, green). In addition, we regenerated the AlphaFoldDB model using the same sequence and confirmed the low RMSD between the AlphaFoldDB and our regenerated structure (*Supplementary Figure 1*). We find that the insertion has significant implications for the ZNT2 structure and locations of the transmembrane and zinc binding domains (*Figures 1C-D*). There were also 221 AlphaFoldDB models where the sequence length was the same, but the sequence differed. In the latter category is Serpin B11 (Q96P1), a non-inhibitory intracellular serpin, with six amino acid substitutions between the two sequences.

**Figure 1:**
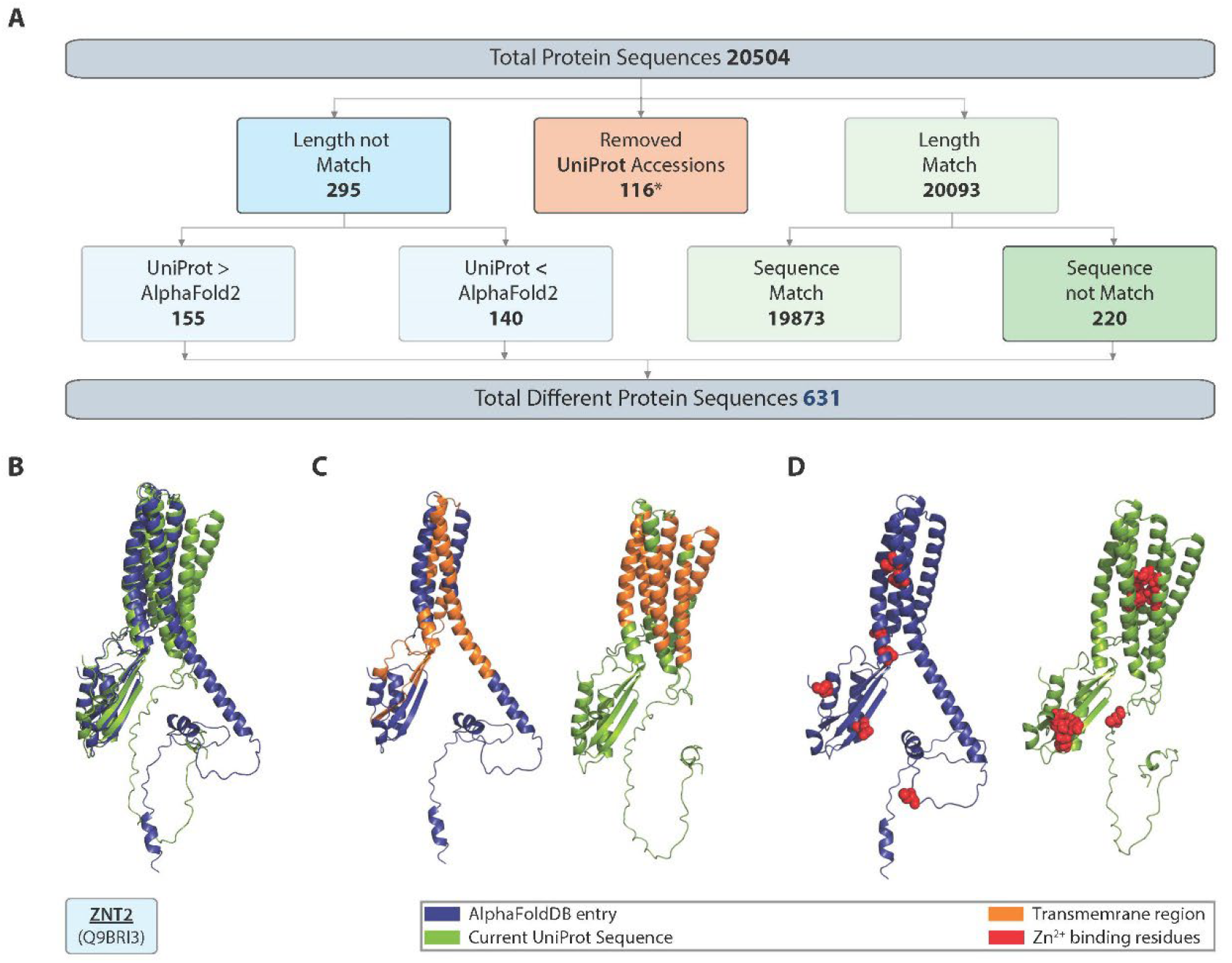
(A) Sequence comparison of human AlphaFold2DB models with corresponding UniProt sequences. Blue rectangles indicate cases where the UniProt sequence length has changed, red rectangles mark accessions that have become redundant, and green rectangles denote sequences where UniProt and AlphaFoldDB match in length. (B) Structure models of proton-coupled zinc antiporter ZNT2 (Q9BRI3). The AlphaFold2-predicted structure downloaded from the AlphaFoldDB (blue) is shown aligned with a new AlphaFold2 model generated from the updated UniProt sequence (green). (C) The transmembrane region is highlighted in orange in both the AlphaFoldDB model (blue) and the UniProt-updated model (green), showing the difference in localisation of these regions. (D) Zinc-binding residues are marked in red on both the AlphaFoldDB model (blue) and the UniProt-updated model (green), highlighting their conserved positions in the protein structure.

To assess the importance of proteins with differing sequences, we found that 406 out of the 631 had high annotation scores (4 and 5), 512 were “reviewed” proteins, and 136 were associated with diseases (as annotated by UniProt). Furthermore, 413 of these proteins were associated with Gene Ontology biological process terms and 418 with molecular function terms, highlighting their potential biological and functional relevance.

We expanded this analysis to include five additional model organisms (Table 1), with mouse showing a very similar trend to human, with 2.5% differing sequences. Interestingly, human and mouse sequences showed greater variation compared to other organisms, probably reflecting the increased depth of research and updates associated with these species. We further investigate this suggestion by looking at the ‘*Date of last sequence modification*’ of UniProt entries across different species. We observe a consistent trend in a higher frequency of changes for human and mouse sequences since 2020 (*Supplementary Figure 2*). On the other hand, *D. melanogaster* and *A. thaliana* had sequence differences of 0.32% and 0.18% respectively with *S. cerevisiae* showing the least difference between AlphaFoldDB and UniProt sequences from all species (0.08%). Remarkedly *D. rerio* shows a 46.68% difference in its sequences, as nearly half of its proteome is absent from the current UniProt version. This is due to a major curation effort that removed thousands of obsolete or redundant entries, reducing the reviewed *D. rerio* proteome to 3,355proteins, of which 93 are not present in the AlphaFold database. Notably, some entries, such as Q802X2 and B7ZC96, have AlphaFold models but are excluded from the downloadable version of AlphaFoldDB, likely due to their annotation as “unreviewed” structures. Reviewing their UniProt history reveals they were only recently marked as reviewed (Q802X2 in July 2024 and B7ZC96 in June 2023), well after the last update of AlphaFoldDB in 2022, illustrating both sequence and annotation aging in the AlphaFoldDB. Excluding all the AlphaFoldDB models no longer corresponding to the updated UniProt database, the total number of differing protein sequences is reduced to 10, resulting in a UniProt AlphaFoldDB difference that aligns with trends observed in other organisms.

**Table 1:**
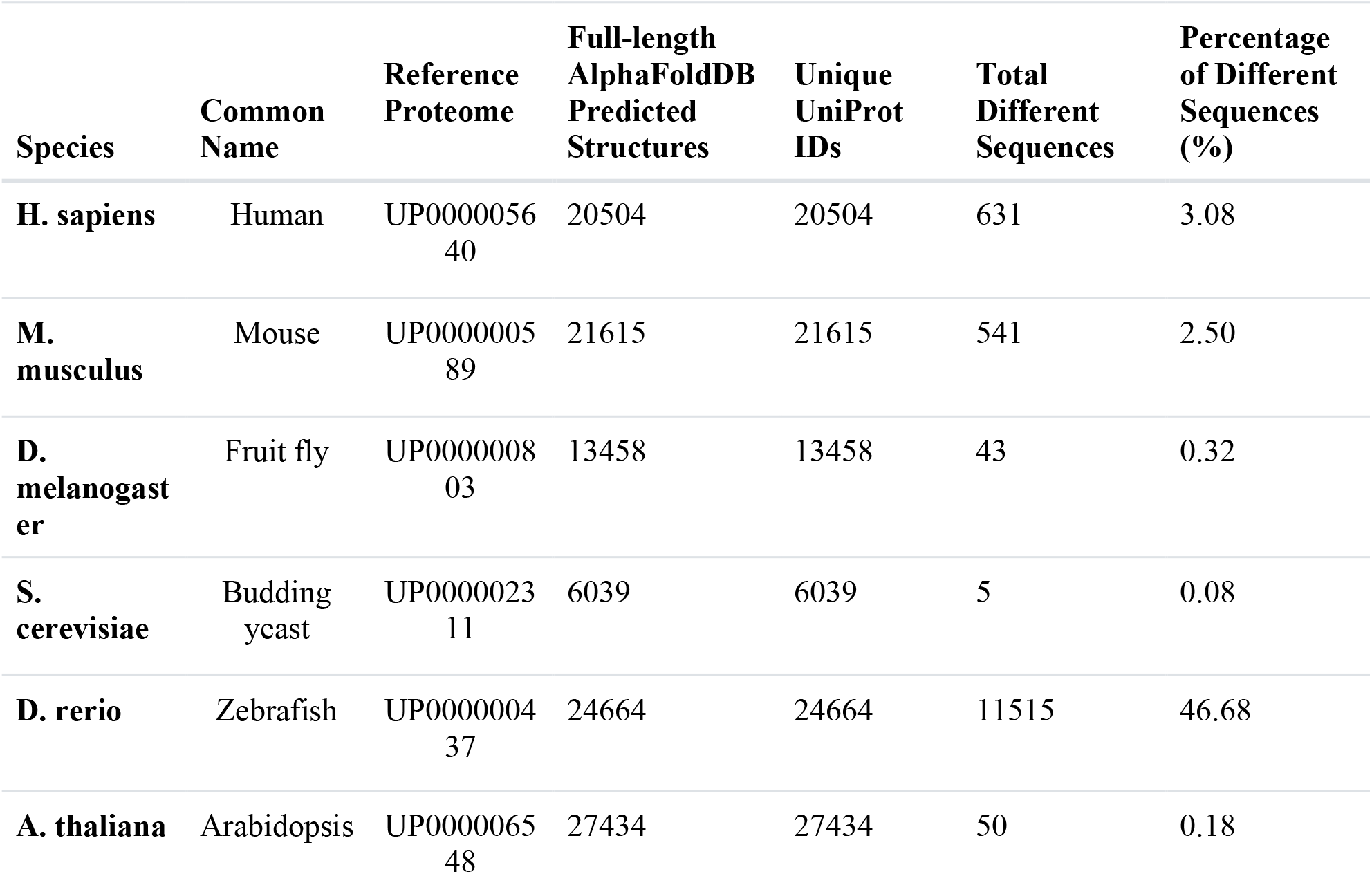
Summary table of the sequence comparison between UniProt and AlphaFold2DB. The percentage of different sequences is calculated on the unique UniProt IDs.

In conclusion, the AlphaFoldDB provides an unprecedented resource for structural biology but its reliance on outdated sequence and annotation data poses challenges across its widespread applications, including structure-based drug design, variant interpretation and functional analysis. These discrepancies highlight the need for regular updates to maintain alignment with UniProt, especially as other groups, such as SwissModel, continue to develop their own structure databases. One way to mitigate these issues is by using UniProt as the primary reference and mapping sequences to the corresponding AlphaFold models through alignment. We stress the importance of routinely checking consistency between UniProt sequences and AlphaFold models, as underscored by UniProt’s and SWISS-MODEL’s new warning flags and recent community efforts to remodel outdated entries with AlphaSync (https://alphasync.stjude.org/) ^11^. However, not all community resources have implemented such warnings yet. Addressing these issues will be critical for maximising the utility of AlphaFold models and ensuring their integration into downstream tools and resources. Similarly, static bioinformatics databases, including AlphaMissense ^12,^ which present precomputed missense variant effect predictions, are also aging. Time flies in bioinformatics.

## Acknowledgements

We thank Harold Powell for helpful discussions

## Supplementary Data (in Excel)

Supplementary Table 1: Table of the sequence analysis between UniProt and AlphaFold2 for *H. sapiens, M. musculus, D. rerio, D. melanogaster, S. cerevisiae* and *A. thaliana*.

## Supplementary Information

Materials, Methods and supplementary figures.

## Data and Code Availability

All data used in this analysis is publicly available. The UniProtKB data (release 2025_03) is available at https://ftp.uniprot.org/pub/databases/uniprot/current_release/. The AlphaFoldDB proteomes are available for download at https://alphafold.com/download#proteomes-section. The analysis is performed using Bash and Python programming languages. The code is available upon request.

## The Aging of the AlphaFold Database - Materials and Methods

### Sequence Analysis & Comparison

First, AlphaFold models were downloaded from https://alphafold.ebi.ac.uk/download ^1^ in accordance with Table 1. Next, UniProt sequences for each AlphaFold model were downloaded directly from UniProt ^2^ along with their respective annotations, including involvement in disease, disruption phenotype, biological process, molecular function Gene Ontology, and date of last sequence modification (version 2025_03, released June 18, 2025). Then, the number of residues in the AlphaFold2/UniProt sequences was compared, and their UniProt Accession Code and Annotation Score was investigated. In addition, the information on the ‘Date of last sequence modification’ was plotted across different sequences using Python and the Plotly library to highlight the frequency of sequence changes across different sequences. The analysis was performed during August 2025.

### AlphaFold2 Model Generation

For the generation of new AlphaFold models, the open-source Colab implementation of AlphaFold2^3^ (simplified version 2.3.2) was used. We regenerated the AlphaFoldDB model using the same outdated sequence (Supplementary Figure 1B, lilac). We modelled the updated UniProt sequence (from version 2025_03) of the entry Q9BRI3 (Figure 1B, green). All structures are modelled using the same default parameters of the monomer AlphaFold2 model and include a relaxation run.

### Structure feature mapping

The transmembrane regions of Q9BRI3 were mapped using DeepTMHMM ^4^, and the Zn2+ binding site annotations were extracted from the UniProt feature viewer. All structure visualisations were performed with the PyMol Molecular Graphics System, Version 3.0 Schrödinger, LLC.

### Structure model alignment and RMSD calculation

Structures are aligned in PyMol. RMSD is computed using PyMol’s rms function. Full atom RMSD between AlphaFoldDB entry and our regenerated AlphaFoldDB model is 15Å. However, when we cut out the long flanking low-pLDDT region (first 65 amino acids of the N-terminus), the resulting full atom RMSD is 0.5Å.

## Supplementary Figures

**Supplementary Figure 1:**
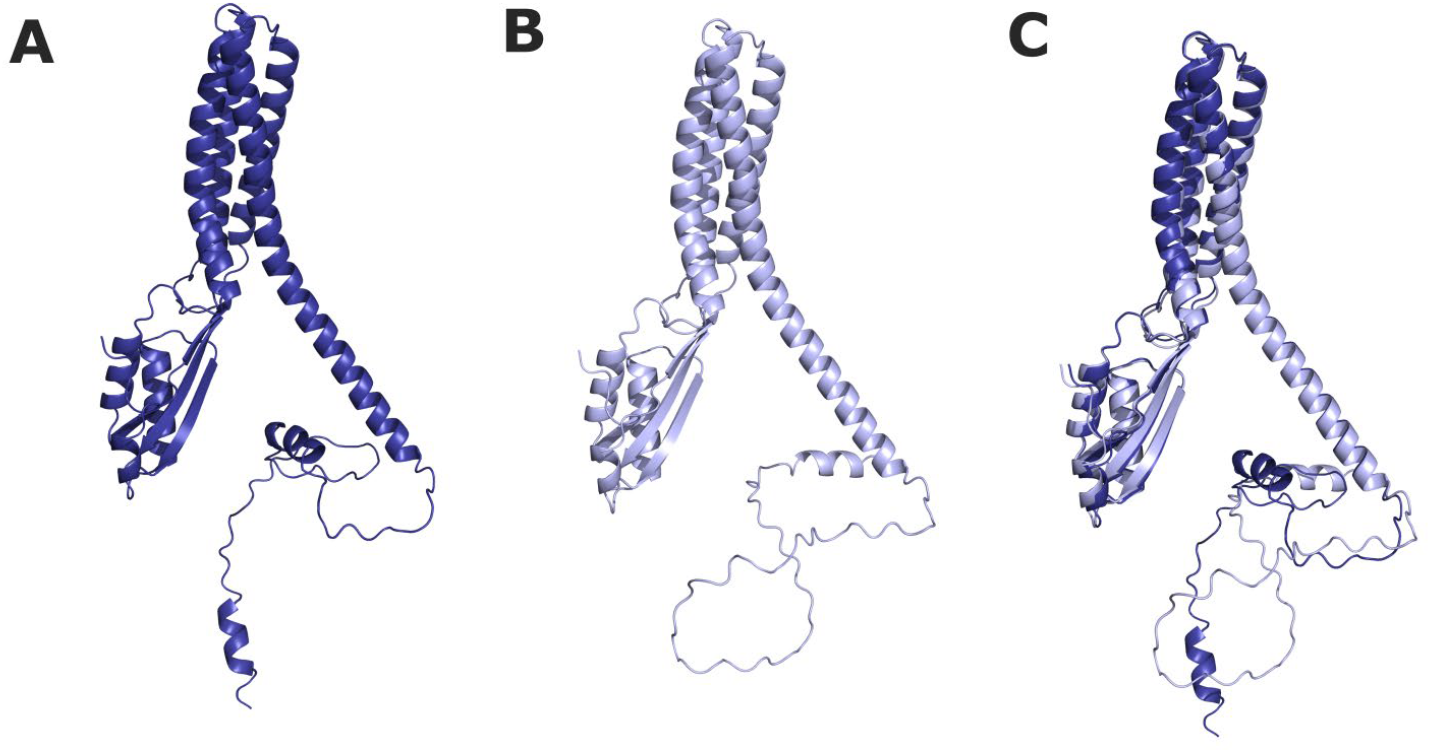
Validation of our AlphaFold2 modelling by regenerating the original AlphaFoldDB model. (A) Original AlphaFoldDB entry for Q9BRI3 (blue); (B) Regenerated AlphaFoldDB model for the same outdated Q9BRI3 sequence (lilac); (C) Alignment between the AlphaFoldDB entry and our regenerated AlphaFoldDB model. The models align well overall, except for the loop in the N-terminus low-pLDDT region. The full-atom RMSD is 15Å, while the RMSD excluding the first 65 amino acids of the N-terminus is 0.5Å.

**Supplementary Figure 2:**
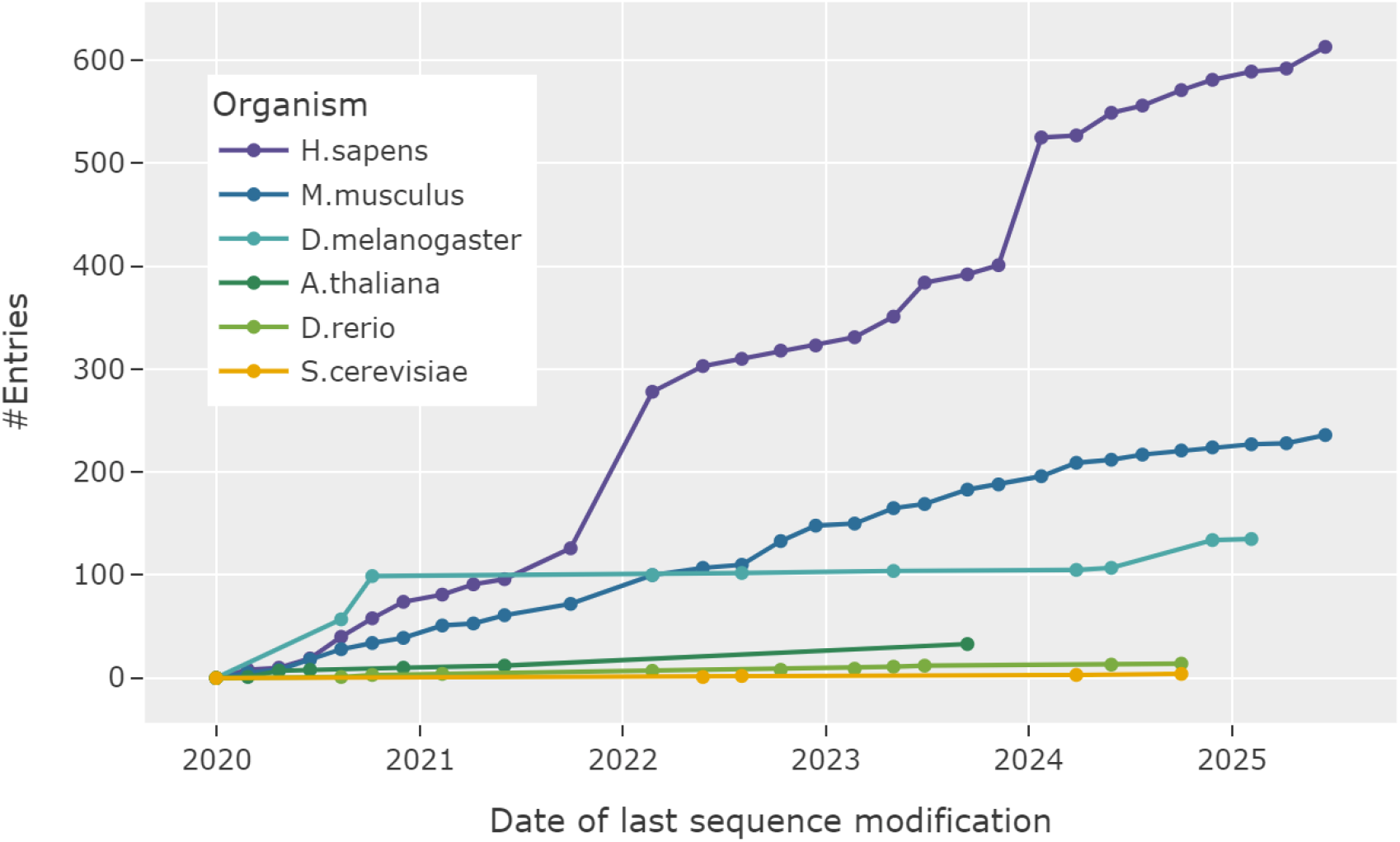
Cumulative number of UniProt entries with sequence changes since January 2020 (across species). The x-axis shows the dates of the last sequence change as reported in the UniProt database. In contrast, the y-axis shows the cumulative number of entries that changed between January 2020 and the date indicated on the x-axis. Colours represent different species. Note that ‘date of last sequence modification’ does not account for the removal of entries from UniProt.

